# Comprehensive investigation of temporal and autism-associated cell type composition-dependent and independent gene expression changes in human brains

**DOI:** 10.1101/065292

**Authors:** Qianhui Yu, Zhisong He

**Affiliations:** CAS Key Laboratory of Computational Biology, CAS-MPG Partner Institute for Computational Biology (PICB), Shanghai Institutes for Biological Sciences (SIBS), Chinese Academy of Sciences (CAS), Shanghai, China; University of Chinese Academy Sciences, Beijing, China

**Author notes:** Correspondence: Zhisong He.

## Abstract

The functions of human brains highly depend on the precise temporal regulation of gene expression, and the temporal brain transcriptome profile across lifespan has been observed. The substantial transcriptome alteration in neural disorders like autism has also been observed and is thought to be important for the pathology. While the cell type composition is known to be variable in brains, it remains unclear how it contributes to the temporal and pathological transcriptome changes in brains. Here, we applied a transcriptome deconvolution procedure to an age series RNA-seq dataset of healthy and autism samples, to quantify the contribution of cell type composition in shaping the temporal and autism pathological transcriptome in human brains. We estimated that composition change was the primary factor of both types of transcriptome changes. On the other hand, genes with substantial composition-independent expression changes were also observed in both cases. Those temporal and autism pathological composition-independent changes, many of which are related to synaptic functions, indicate the important intracellular regulatory changes in human brains in both processes.

## Introduction

The development and aging of human brains are complex processes, which are shaped by anatomical and molecular changes^1-4^. Autism, as a common neurodevelopmental disorder, disrupts the critical developmental processes and results in the disruption of cognitive functions^5-7^. With the emergence of high-throughput measurement of different molecules, dozens of studies have been conducted to characterize age-related molecular changes in human brains, especially at the transcriptome level^8-11^. Meanwhile, several studies have also been conducted in order to investigate the transcriptome alteration in autism^12-14^.

The human brain, however, is a highly complex and heterogeneous organ comprised of numerous different cell types, including neurons and multiple classes of non-neuronal glial cells – such as astrocytes, oligodendrocytes, oligodendrocyte precursor cells and microglia – as well as vascular, such as endothelial, cells. Each of those cell types expresses a distinct set of genes^15^ and plays a unique and essential role in the development and functions of the brain^16^. Different cell types are also known to show different spatial-temporal distributions. Such complexity thus raises the unanswered questions: how much of the age-related molecular change in human brains, or specifically the age-related transcriptome change, is the direct consequence of the cell type composition change? If composition changes contribute to age-related expression changes, what’s the biological meaning of the rest? Similar questions about the molecular changes in neural disorders, including autism, also remain opened.

In order to comprehensively answer these questions, an accurate estimation of cell type composition in healthy and autistic human brains is required. Although experiments including stained cell counting^17-20^ and large scale single-cell RNA-seq^21,22^ have the potential to provide these data, these methods are either too labor-intensive or costly at present. Thus, the computational method of inversing sample heterogeneity, *i.e.* deconvolution, is one of the best alternative solutions to estimate the mixing percentage of different cell types^23-25^. The application of transcriptome deconvolution to human brain was long limited by the lack of comprehensive transcriptome data for the varied cell types in human brains. The recent emergence of the human brain single cell RNA-seq data, covering all the main cell types in the human brain^21^, now renders this approach more feasible.

In this study, we used a two-step deconvolution procedure to estimate the cell type composition changes in the human brain after birth, as well as to quantitatively dissect the human brain transcriptome profiles across lifespan into composition-dependent and composition-independent components. The estimated composition changes matched well with the previous observations. The delineation of the composition-dependent component from gene expression suggested that cell-type composition explained about 25% of the total expression temporal variance, and greatly contributes to the age-related expression pattern in brain. Meanwhile, although to a lesser extent, the composition-independent component also significantly contributes to age-related expression pattern. In addition, we used the framework to analyze the contribution of cell-type composition alteration in the brain transcriptome alteration of autism patients. While composition-dependent component, especially the decreased proportion of RNA contributed by neurons, appeared to be the main power driving the difference, one group of synaptic genes with altered composition-independent component presented strong enrichment of autism-associated genetic variants, implying the active regulatory disruption of synaptic functions in autism.

## Results

### Cell type composition changes in brains across the human lifespan

To investigate the temporal cell type composition changes across lifespan in human brains, we applied the deconvolution procedure to the age series RNA-seq data set of the human prefrontal cortex (PFC) consisting of 40 postnatal human brain samples aged from two-day old to 61.5 year-old^11^. A published human brain single cell RNA-seq data^21^ was used to obtained the gene expression information of the eight main cell types in human brains, quantifying the expression level of 14,054 protein-coding genes. CIBERSORT^24^ was used to select genes distinguishing cell types, resulting in 904 genes (referred as CIBERSORT-selected markers, Supplementary Dataset S1). Two well-known transcriptome deconvolution algorithms, CIBERSORT^24^ and quadratic programming (QP), were applied based on the CIBERSORT-selected markers.

In addition, diffusion ratio (DR) based deconvolution was also used based on the 1,491 cell type signature genes. This deconvolution algorithm was based on the assumption that the expression of a cell type signature gene in the bulk tissue can be seen as its expression in the cell type scaled by the cell type’s mixing proportion. It is similar to the recent deconvolution algorithm MCP-counter^26^ but simplified (see Methods). The cell type signature genes were identified by requiring at least ten-fold higher expression level in one cell type comparing to any of the remaining, resulting in 319 signature genes for astrocytes, 288 for endothelial cells, 224 for microglia, 99 for oligodendrocytes, 71 for oligodendrocyte progenitor cells (OPC), 166 for mature neurons, 92 for immature neurons, and 232 for replicating neurons (Supplementary Dataset S1).

The three deconvolution algorithms resulted in similar composition patterns across the postnatal lifespan (Figure 1 and Supplementary Figure 1). The composition pattern of DR algorithm was robust to confounding factors, e.g. neuron heterogeneity, neuron ageing and sex difference (PCC=0.99, Supplementary Figure 2), as well as cell variability in the single cell RNA-seq data (median PCC=0.94, Supplementary Figure 3). The estimated composition changes were consistent with previous studies, e.g. the elimination of immature neurons soon after birth accompanied with the increase of mature neurons^27^, as well as the decrease of OPC (linear regression slope test, *P*=0.0016) accompanied by the trend of increase of oligodendrocytes (linear regression slope test, *P*=0.21) which may be due to the myelination process^28^. The remaining cell types including astrocytes, endothelial cells and microglia did not show significant changes across the lifespan.

**Figure 1.**
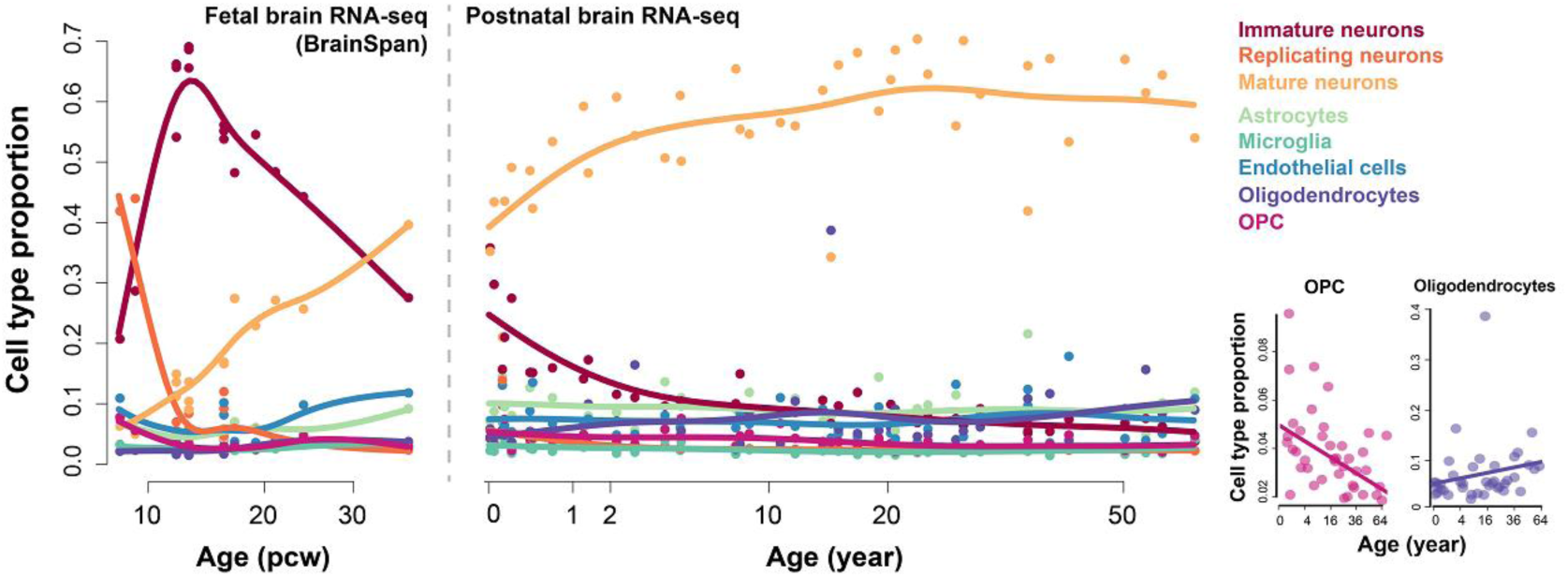
The cell type compositions across lifespan in human brains, estimated using the DR-based deconvolution. The dots show the estimated proportions of samples, and the curves show the spline interpolation results. The left panel shows the cell type composition in human fetal brains, based on the human embryonic developmental brain RNA-seq data from BrainSpan; the right panel shows the cell type composition in human postnatal brains, based on the human postnatal brain data set. The bottom-right panels show proportions of OPC and oligodendrocytes in the human postnatal samples.

Nevertheless, the three deconvolution algorithms provided slightly different results. Therefore, two complementary measurements, namely the similarity measurement and discrimination measurement (see Methods), were used to compare the three estimated composition patterns. According to Pearson’s correlation coefficient (PCC) and root-mean-square deviation (RMSD), both of which were similarity measurements, CIBERSORT performed the best among the three algorithms, while DR performed slightly better than QP (Supplementary Figure 4). On the other hand, the discrimination measurement (see Methods), suggested that the cell type gene expression identities were reconstructed in the most discriminating way by DR, while CIBERSORT performed better than QP (Supplementary Figure 4).

In addition to the postnatal brain transcriptome, we further applied the three algorithms to the human embryonic developmental brain RNA-seq data obtained from BrainSpan database (Figure 1 and Supplementary Figure 1). DR-estimated composition pattern successfully recapitulated the domination of replicating neurons dominate before 12 post conception weeks (pcw) and its dramatic elimination after that. This observation, coupled with the increase of immature neurons, matched well with the neuronal proliferation that occurred between four pcw and 12 pcw^29^. Moreover, compositions estimated by DR algorithm, especially those of immature and mature neurons, presented a successive pattern with the postnatal composition changes estimated above, which further indicated the reliability and robustness of the composition estimation. On the other hand, CIBERSORT, although performed better than both DR and QP deconvolution according to the similarity measurement, failed to reproduce those scenarios.

### Composition change is the primary power shaping the human brain postnatal temporal transcriptome

To comprehensively study the contribution of cell type composition changes in the human brain temporal transcriptome, we dissected the bulk gene expression into the composition-dependent and composition-independent components (Figure 2), based on the composition pattern estimated by DR algorithm (see Methods). The composition-dependent component represents gene expression changes related to cell type proportion difference, while the composition-independent component represents changes which are independent of composition changes and may be related to change of intra-cell type heterogeneity or molecular features, such as activity changes of particular pathways or biological processes in certain cell types (Figure 2).

**Figure 2.**
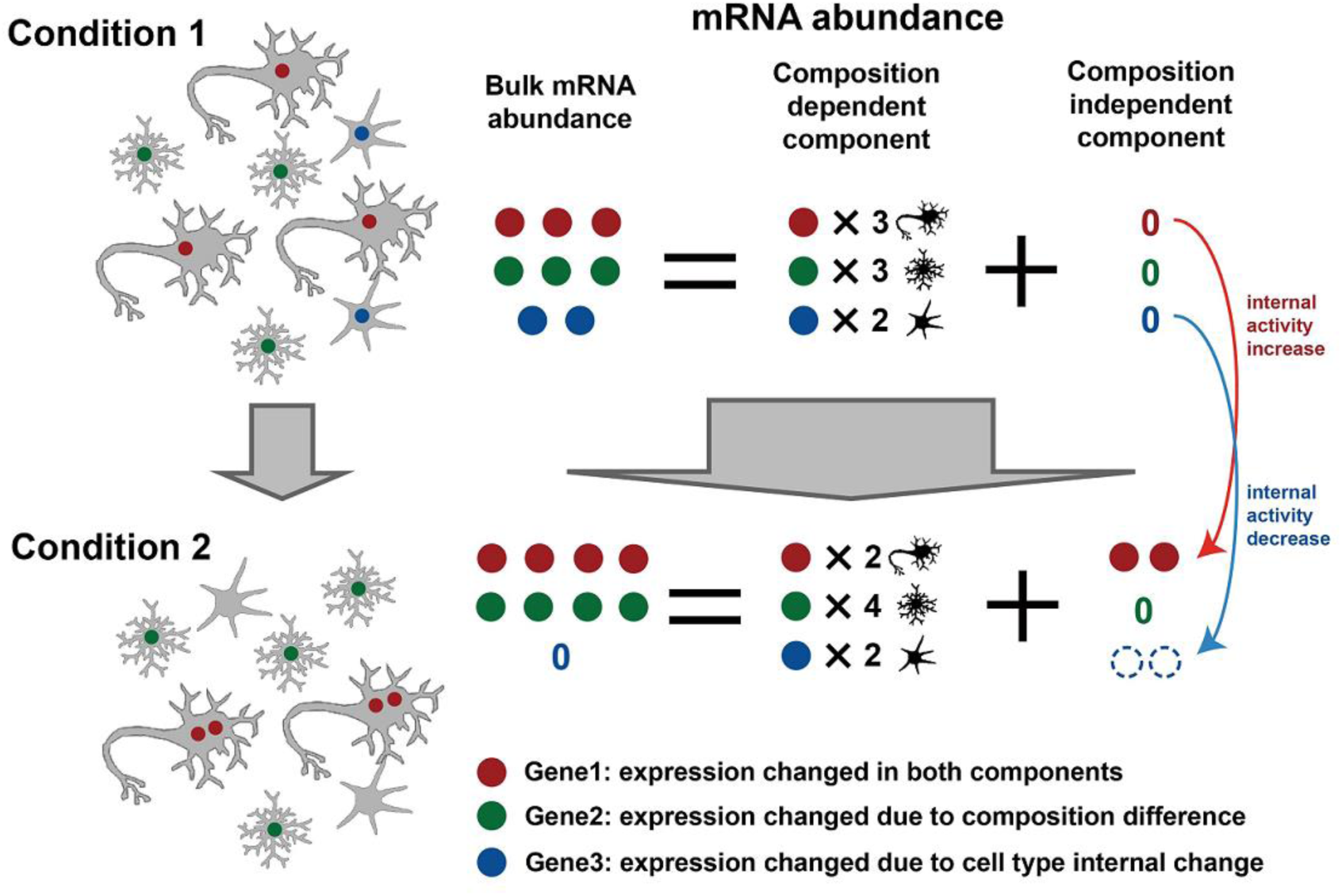
Schematic representation of gene expression dissection into composition-dependent and composition-independent components. Circles with different colors represent mRNA products of different genes. Dashed hollow circles in composition-independent components represent “negative” expressions, *i.e.* the observed expression level in bulk tissue is lower than the expected expression level when composition difference is assumed to be the only variable (*i.e.* composition-dependent component).

We then estimated the contributions of the two components to the overall gene expression variance. On average, the composition-dependent component, which represents the upper limit of transcriptome variance caused by variation of cell type composition in human brains, explained 26.4% of the total variance in the human postnatal age series data (Figure 3a). Focusing on the 5,119 genes with age-related expression (referred as age-related expressed genes, age test Benjamini-Hochberg (BH) corrected *P*<0.05), the composition-dependent component contributed 34.7% of the total variance which was significantly higher than the other genes (Figure 3a, permutation test, *P*<0.001).

**Figure 3.**
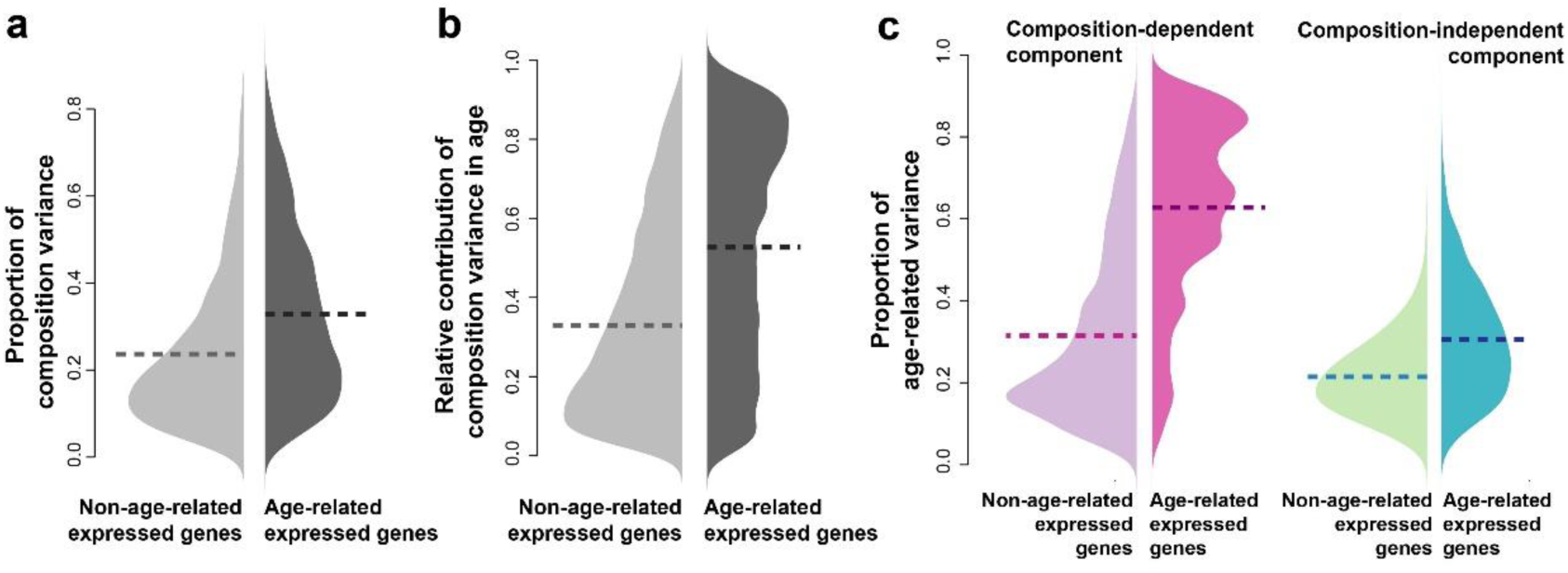
The cell type composition change was the primary driving force of the age-related expression. (a) The proportion of gene expression variance explained by cell type composition (composition-related variance) in the human postnatal brain data set. Light grey – genes without age-related expression; dark grey – genes with age-related expression. The horizontal dash lines show the mean proportion of composition-related variance. (b) The relative contribution of the composition-dependent component to the variance explained by age (age-explained variance), measured as ratio of age-explained variance in the composition-dependent component to the sum of age-explained variance in both components. Light grey – genes without age-related expression; dark grey – genes with age-related expression. The horizontal dash lines show the mean proportion of composition-related variance. (c) The proportion of variance explained by ages (age-related variance) in each of the two components of expression: pink – the composition-dependent component; green – the composition-independent component. The light colors represent the proportions of age-related variance in genes without age-related expression changes; the dark colors represent the proportions of age-related variance in genes with age-related expression changes. The horizontal dash lines show the means of age-related variance proportions.

To further investigate the roles of the composition-dependent and independent components in shaping the temporal gene expression patterns, we calculated the variances explained by age (age-explained variance) for each of the two components separately. Interestingly, the relative contribution of age-explained variance from the composition component (55.8%) was much larger than the proportion of composition variance among total variance (34.7%) for age-related expressed genes, while the difference was much smaller for the non-age-related expressed genes (27.8% vs. 22.1%) (Figure 3b). Altogether, these observations implied that the change of the composition-dependent component, *i.e.* the cell-type composition changes, was the main power shaping the observed temporal expression in human brains.

Additionally, the proportion of age-explained variance in the composition-dependent component was dramatically higher for the age-related expressed genes than for other genes (median=66.4%, permutation test, *P*<0.001, Figure 3c). Meanwhile for the same genes, a moderate but significant increase of age-related variance proportion was observed for the composition-independent component (median=28.9%, permutation test, *P*<0.001, Figure 3c). These observations not only point to the significance of composition-dependent changes, but also to that of the composition-independent changes, some of which may represent the changes of molecular features in one or several cell types, also participate in shaping the temporal transcriptome in human brains. It is worth to mention, that all the above observations were reproducible based on CIBERSORT-based composition pattern (Supplementary Figure 5), suggesting that it is not an artifact of the DR-based deconvolution procedure.

### The age-related changes in the composition-independent components are important and well regulated

To better understand the biological significance of the age-related changes in the composition-dependent and composition-independent component explicitly, we applied age tests to each of the two components (see Methods), to identify genes with significant age-related changes in either component. 8,156 and 1,455 genes were found with age-related changes in the composition-dependent and independent component, respectively (Figure 4a, Supplementary Dataset S2; ANCOVA, BH corrected *P*<0.05). Both of the two gene sets were largely overlapped with the age-related expressed genes (Fisher’s exact test, *P*<0.0001). However, no significant overlap was observed between them (Fisher’s exact test, *P*=0.216, odds ratio=0.932), implying the independent contribution of the two components to the temporal transcriptome in human brains. CIBERSORT-based composition pattern provided similar estimation, resulting in 6,671 and 1,129 genes with age-related changes in the composition-dependent and independent component, respectively. Majority of those genes were overlapping with genes identified based on DR-based estimation (Supplementary Figure 6).

**Figure 4.**
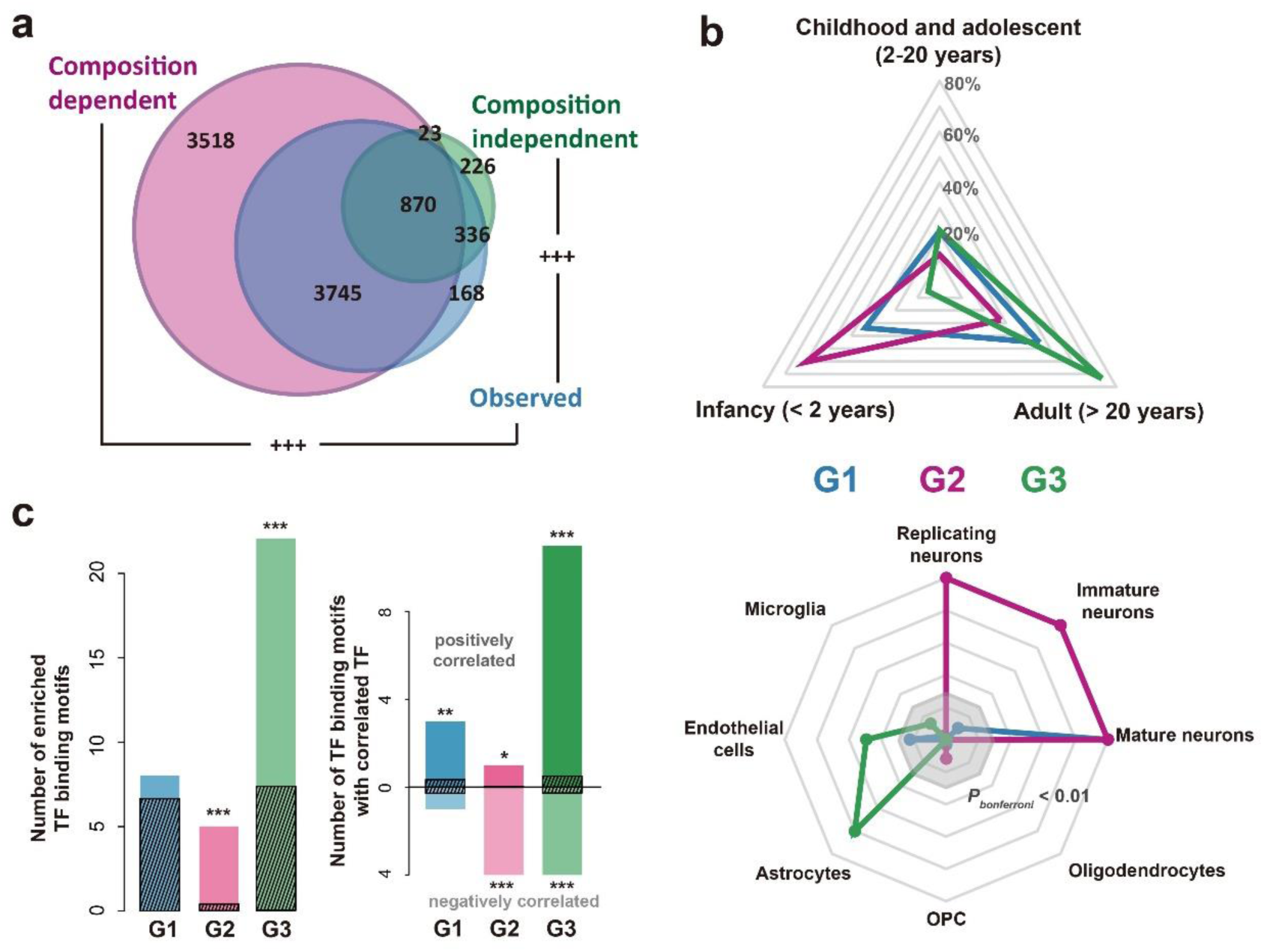
Age-related changes happened in the composition-dependent components and composition-independent components. (a) The number of genes with age-related changes in each component or the observed expression level, based on the human postnatal brain data set (blue – observed expression level; pink – composition-dependent component; green – composition-independent component). (b) The expression properties of genes with age-related changes in the composition-dependent or composition-independent components. G1 – genes with age-related changes in both components; G2 – genes with age-related changes only in the composition-dependent component; G3 – genes with age-related changes only in the composition-independent component. Top: proportion of genes with highest expression level at each of the three lifespan stages. Bottom: expression enrichment in each of the eight cell types for the three groups of genes, represented as –log10(*P*), where *P* being the p-value of one-sided Wilcoxon’s rank test of log10-transformed fold change from the particular cell type to the remaining cell types, between each of the three groups of genes and all the expressed protein-coding genes. The grey octagonal boxes represent –log10(*P*) equaling to values from ten (the outermost box) to two (the inner most box). Strong expression enrichment with –log10(*P*)>10 was presented as ten. (c) Regulation of genes with age-related changes in either component by transcription factors (TFs). (Left) the number of TF binding motifs enriched among genes within each group. The dark streaked bars represent the mean number of enriched TF binding sites expected by chance, calculated by 1000 random assignment of the expressed genes into the three groups. (Right) the number of TF binding motifs with its representative TF correlated with the targets (correlated TF binding motifs) in the same group. The dark streaked bars represent the mean number of correlated TF binding motifs expected by chance, calculated by 1000 random assignment of the expressed genes into the three groups. The asterisks show significance of the numbers (* *P*<0.05, ** *P*<0.01, *** *P*<0.001, Bonferroni corrected).

We further grouped the 8,719 genes with age-related changes in at least one of the two components into three categories: G1 – genes with age-related changes in both components; G2 – genes with age-related changes only in the composition-dependent components; and G3 – genes with age-related changes only in the composition-independent components. Interestingly, these three groups of genes showed distinct temporal expression patterns (Figure 4b). G1 and G2 genes, and especially the latter ones, showed higher expression levels in early postnatal development stages, while G3 genes were highly expressed in adult stages. Different groups of genes were also enriched in different cell types: G1 – mature neuron, G2 – immature neuron, replicating neuron and mature neuron, and G3 – astrocyte and endothelial cells (Figure 4b).

More interestingly, the three groups of genes showed distinctive functional enrichments (DAVID ^30^, Supplementary Table S1). In brief, G1 genes were enriched for synapse and translation-related functions. G2 genes were enriched in transcription regulation, protein degradation, and cell cycle. Lastly, G3 genes were significantly involved in extracellular regions and metabolism. These results indicated the distinct biological significance of the age-related composition-dependent and independent changes.

The expression trajectories of these three groups of genes may have been modulated by different regulatory mechanisms, such as transcription factors (TFs). To test this, we estimated the enrichment of TF binding motifs in the promoter regions of genes in each category. We observed significant excess of enriched TF binding motifs (hypergeometric test, *P*<0.1) in the G2 and G3 genes (permutation test, *P*<0.001) (Figure 4c). In addition, the representative TFs’ expression patterns of the enriched TF binding motifs showed significantly better correlation (Wilcoxon test, *P*<0.05) with their targets in the respective groups than expected by chance (permutation test, *P*<0.001) (Figure 4c). TFs with TF binding motifs enriched in G2 genes, e.g. CUX1 and E2F1, were mostly negatively correlated with their targets and had been shown to be relevant to cell migration, cell cycles and neuronal development and maturation ^31-33^.On the other hand, most of the TFs with binding motifs enriched in G3 genes, e.g. SMAD3, SREBF1 and NR2F2, were positively correlated with their G3 target genes, many of which had been reported participating in signal transduction and metabolism of astrocytes ^34-36^.

### Significance of composition-independent component in autism pathogenesis

Autism is a common neurodevelopmental disorder. Previous transcriptomic studies have suggested dramatic changes of gene expression ^12-14^. For instance, using the same human PFC postnatal age series RNA-seq data set as described above, coupled with RNA-seq data of 34 PFC samples of autistic patients, Liu et al. identified 1,775 genes with differential expression (DE) between autism cases and unaffected controls 14. It is worthwhile to estimate the contribution of cell type composition as well as to identify the composition-independent alteration which may represent the active regulatory alternation in autism patients.

We applied CIBERSORT, QP and DR algorithms to estimate composition patterns in the autistic samples. The three algorithms provided similar results. The composition differences by DR deconvolution were robust to confounding factor (PCC=0.90, Supplementary Figure 2), as well as cell variability in the single cell RNA-seq data (median PCC=0.84, Supplementary Figure 3). Interestingly, results of the three methods all showed significant smaller proportion of neurons as well as larger proportions of non-neuron cells (Figure 5a and Supplementary Figure 7). The amplitudes of composition changes were significantly correlated among the estimations based on the three algorithms (Supplementary Figure 7), eliminating the possibility of algorithm artifact. The general trend of decreased expression levels of neuronal signature genes in autistic brains, as well as the increased expression levels of astrocytes, microglia and endothelial cells signature genes, were also observed in another microarray dataset^13^ (Supplementary Figure 8). The increase amplitudes of different non-neuron cell types were proportional to their estimated proportion in healthy samples (Figure 5b), suggesting that they were likely to be the passive consequence of neuron proportion decrease.

**Figure 5.**
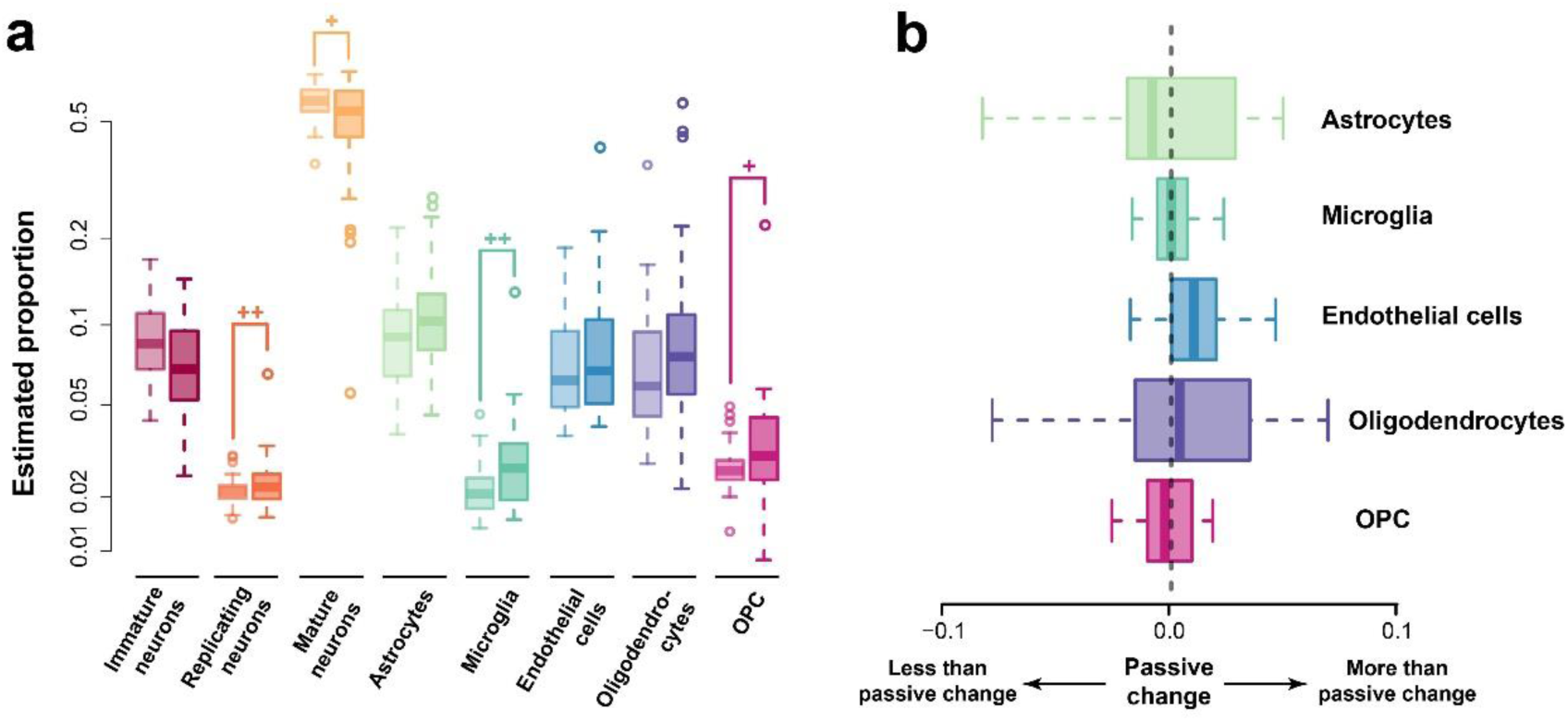
Alteration of cell type composition in autism. (a) The estimated proportions of each cell types in autism case samples and healthy control samples with ages at the same age range. Each pair of boxes with the same color show the proportions of one cell type, with the left box represents the proportions in healthy samples, and the right box represents the proportions in autism samples. The plus signs mark significant difference according to Wilcoxon’s test (+ – *P*<0.1; ++ – *P*<0.05). (b) The disparity of observed altered non-neuron cell type compositions and the predicted altered compositions with the assumption of passive change of non-neuron cell type composition. Each box with a different color showed the observed altered proportion of one non-neuron cell type subtracted by the predicted altered proportion of the cell type based on the altered proportion of neurons scaled by the relative proportions of the non-neuron cell type in healthy samples.

To further investigate the contribution of composition differences in the autistic brain transcriptome alteration, we dissected the bulk gene expressions of all the PFC samples, including the 34 autism cases and 30 healthy controls within the same sample age range, into the composition-dependent and composition-independent components, based on the DR-based composition pattern. Natural spline based ANCOVA (Methods) was applied to each of the two components to identify genes with DE between autism cases and healthy controls. Among the 14,032 detected genes, 74% of them (10,358) showed DE in their composition-dependent components (BH corrected *P*<0.2), while less than 5% of the genes (644) showed DE in composition-independent components under the same criteria (Supplementary Dataset S3). The two numbers based on CIBERSORT-based composition pattern were 8,030 and 275, with majority overlapping with the DR-based ones (Supplementary Figure 6). Both the two sets of genes were significantly overlapping with the 1,775 DE genes identified based on bulk expression (referred as bulk DE genes, Figure 6a, Fisher’s exact test, *P*<0.0001), suggesting that both of them were important contributors in transcriptome alteration in autism brains.

**Figure 6.**
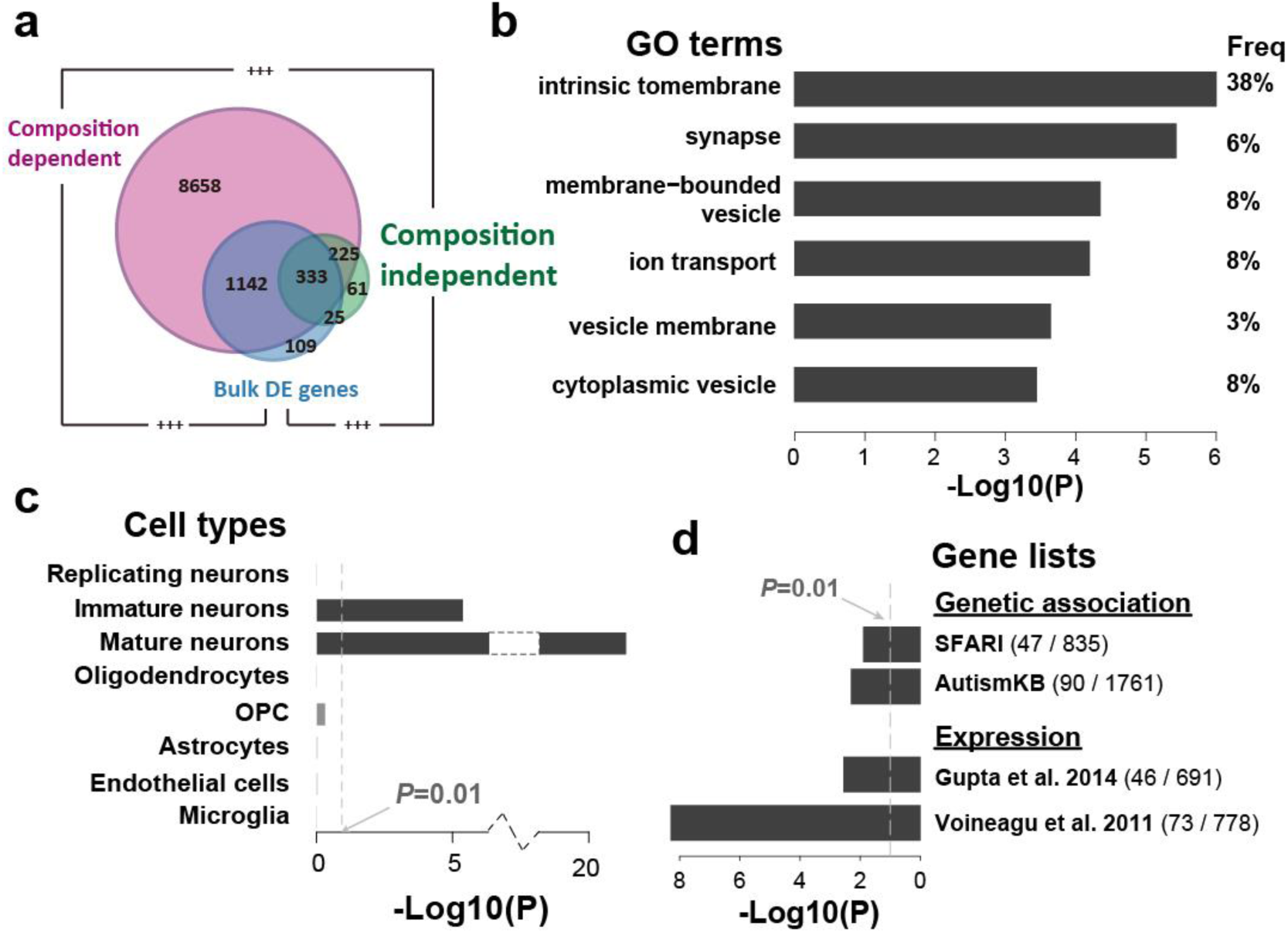
Composition-dependent and independent transcriptome alteration in autism. (a) The number of genes with autism alterations in each component or the bulk expression levels (blue – bulk expression levels; pink – composition-dependent component; green – composition-independent component). The plus signs mark the significance of overlap (+++ - Fisher’s exact test *P*<0.0001). (b-d) Functional characterizations of genes with autism alterations in the composition-independent components, including GO enrichments (b), cell type expression enrichments (c) and enrichments of autism-associated genes (d). The bars show the log10-transformed *P*-values. The values within brackets in the button right panel show the numbers of all associated genes (right) and the sizes of the overlap with genes with autism alterations in their composition-independent components (left).

Although majority of the transcriptome alterations in autism brains were related to the cellular composition changes, the alterations in the composition-independent components of brain transcriptome were of particular interest. They might represent the active regulatory disruptions in particular brain cells. The functional enrichments (DAVID ^30^) suggested that they were enriched for membrane and synaptic proteins (Figure 6b, Supplementary Table S2). Consistently, those genes showed significant expression enrichment in mature and immature neurons (Figure 6c). More interestingly, the genes with significant alterations in their composition-independent components, but not those with significant alterations only in their composition-dependent components, contained a significant excess of genes with genetic variants associated with autism, based on analysis of genes extracted from SFARI AutDB ^37^ (Fisher’s exact test, odds ratio=1.52, *P*=0.009, Figure 6d), as well as genes from AutismKB database ^38^ (Fisher’s exact test, odds ratio=1.37, *P*=0.008, Figure 6d). Furthermore, those genes were also identified as differentially expressed genes in autistic brains in previous studies ^12,13^ (Fisher’s exact test, *P*<0.005, Figure 6d). It is also worth to note, that 40 microRNA (miRNA) families showed enrichment of predicted binding at the 3’UTRs of those genes (TargetScan v7.1 ^39^; hypergeometric test, *P*<0.05), which was significantly more than by chance (permutation test, *P*<0.01, *FDR*=0.65). Some of the enriched miRNAs, including has-miR-363-3p which was also reported to be dysregulated in Alzheimer’s disease ^40^, have been reported as differentially expressed in autistic brains ^41^. All the above observations suggested that the disruptions of neuronal transcriptional regulations in autistic brains, which were likely to be partially caused by the dysregulations of certain neuronal miRNAs, may result in pathological changes in synapses and largely contribute to autism pathology.

### Discussions

In this study, we attempted to investigate the role of changes related or independent to cell type composition in human brain development and pathogenesis of autism using computational approaches. With the gene expression of eight main cell types in human brain estimated using the human brain single cell RNA-seq data^21^, we applied a two-step transcriptome deconvolution procedure to a human brain postnatal age series RNA-seq data set^11^ and an associated autism brain RNA-seq data set^14^. Based on the estimated composition patterns, gene expression was dissected into its composition-dependent and composition-independent component, allowing further investigations on them separately.

The transcriptome deconvolution in healthy samples resulted in temporal composition patterns consistent with the prior studies. On the other hand, we also notice that our estimated proportion of total neurons in the adult samples reached around 60-70%, which is much higher than the estimation based on cell counting^42,43^ or DNA methylation deconvolution^44^. This may be explained by the fact, that the estimated composition of transcriptome deconvolution actually reflects the proportion of RNAs contributed by each cell type instead of the proportion of cell numbers. In fact, the RNA content in neurons is reported to be close-to-two-fold as much as in glia cells^45^.

Based on the variance analysis, we regard the composition change to be the main source of the age-related expression change in human brain. This is a consistent conclusion drawn by previous studies based on DNA methylation^44^. The proportion of variance explained by cell types, on the other hand, is smaller: only about 30% for the age-related expressed genes, while it is around 50% for the DNA methylation. Although the effect of noises during sampling and measurement could not be ruled out, such observation implies additional regulatory signal in age-related manner across lifespan, which is independent from DNA methylation and may be an interesting focus for further study.

Interestingly, genes with age-related changes in the composition-dependent components show high expression level in the infant stages, *i.e.* < 2 years old, with significantly enriched expression in the three types of neurons. This suggests that although some other cell types such as oligodendrocytes and OPCs also presented temporal proportion changes, most of the composition-dependent changes are due to the transition from immature neurons to mature neurons, *i.e.* the neuron maturation process. It therefore implies that the neuron maturation is the primary factor creating the age-related expression, especially during the infant stage.

On the other hand, the age-related changes to the composition-independent component is appealing. Unlike the genes with age-related composition-dependent changes which mostly are highly expressed in the early postnatal development, genes with age-related composition-independent changes tend to have higher expression in adults. What’s more, these composition-independent changes are significantly related to either synapse in neurons (G1), or extra-cellular regions and signal peptides in astrocytes and endothelial cells (G3), both of which are relevant to cell-to-cell communications. This is unlikely to be a coincidence. As an information processing system, communications are critically important for human brains^46,47^. Our results suggest that the complexity growth of the communication system not only depends on the increased number of computational units, *i.e.* neurons, but also greatly relies on the enhanced inter-cellular communications which are independent from the cell type composition changes across lifespan. Such communications may not limit to the synaptic connections between neurons, but also include the neuron-glia and glia-glia communications which are also critical to the neuronal network functions^48^.

In autistic brains, the contradictory statements of unchanged density of both neuronal and glial cell pools^49^ and even more neuron numbers^50^ has been reported. Therefore, it is a surprising observation that the estimated neuron proportions in autistic brains are significantly smaller than that in healthy brains. While the influence of unknown technical issue cannot be ruled out, this result implies the possibility that the general transcriptional activity of neurons is inhibited in autism. In such case, the total mRNA content in neurons reduces, therefore resulting in smaller contribution of neuronal mRNAs to the bulk mRNA pool. This assumption is supported by the depletion of neuronal marker MAP2 in autistic brains^49^. However, further experimental measurement is also awaited for validation.

Considering that neuroinflammation is thought to play important roles in autism pathology^51,52^, it is also unexpected to find no evidence suggesting general activation of microglia, which are the macrophage cells participating neuroinflammation in the central nervous system. Although the increase of microglia proportion is observed, no significant excess of increase is observed when comparing to other glial cell types after normalizing to their relative proportion in healthy brains. Nevertheless, our results did not rule out the importance of neuroinflammation in autism, considering that the strongest activation of microglia in autism patients happens in cerebellum and white matters^52,53^. Although higher microglia density was also reported in autism dorsolateral PFC ^54^, they may not be fully activated, resulting in insufficient power in our analysis to capture it. More investigation is required to comprehensively study the role of PFC microglia in autism brains.

While our results suggest that majority of the transcriptome alteration detected in PFC of autism patients are passive alteration due to the changes of relative mRNA contribution from cell types, a group of genes attracts most of the focus. Those 644 genes, enriched for membrane protein genes and synaptic functions in neurons, represent the transcriptome alterations which are at least partially independent to the composition changes. This is an interesting observation, suggesting the autism pathological alteration in the neuronal regulatory network which regulates synaptic functions. This linkage is further supported by their enriched genetic association with autism. It is also concordant with previous studies, which report the abnormalities of synaptic functions as well as neuronal dendrites and spines in autistic humans and autistic mouse models^55,56^. More comprehensive investigations are awaited for the roles of synaptic disruptions in autism, which may lead to effective pharmacological treatment in the future.

Although our framework paves a way for more exhaustive analyses regarding the contribution of cell type composition to the transcriptome changes, we are well aware of the limitations of our method. For instance, our analysis failed to squarely pinpoint the one or several cell types with the greatest influence on either of the two components, though cell-type enrichment analysis was capable to give a glance. Our analysis greatly relied on the accurate transcriptome measurement of all or at least close to all of the cell types in the bulk tissue, which limits its applications. The experimental validation is relatively difficult. Nevertheless, the fast-developing single-cell technology, especially the single-cell RNA-seq, should be able to comprehensively, simultaneously and separately quantify changes happened in cell type composition, cell type RNA content and cell type molecular profiles. Indeed, the single cell RNA-seq technology has been widely used to describe cell type compositions in different organs including brains^21,22,57^, as well as to characterize the molecular processes driving neurogenesis and somatic reprogramming to neurons ^58,59^. The further application of this technology to characterize human brain development, ageing, and pathology of neural disorders including autism would be sufficient to verify our observation and of great value to comprehensively understand those important processes.

If nothing else, we hope that our attempt to dissect expression into composition-dependent and composition-independent components will inspire further studies to elucidate transcriptome changes in a more comprehensive manner.

## Methods

### Data

The human brain single cell RNA-seq data was retrieved from SRA (SRP057196). The human postnatal age-series brain RNA-seq data with 40 samples was retrieved from GEO (GSE51264). All the RNA-seq reads were mapped to the human genome hg38 with STAR 2.3.0e using the default parameters^60^. The number of reads covering the exonic regions of each protein-coding gene annotated in GENCODE v21 was counted and normalized using the R package DESeq261 for each data set separately. The autistic brain RNA-seq data retrieved from GEO (GSE59288), were processed in the same way. The pre-calculated RPKM of the fetal human brain samples were downloaded from BrainSpan (http://www.brainspan.org/static/download.html).

### Deconvolution for cell-type composition

Three different strategies were used for deconvolution for cell-type composition. The first method, quadratic programming (QP) based deconvolution, was to model the gene expression of each cell type signature genes in the bulk tissue sample as a linear combination of its expression in each cell type according to the cell type mixing proportion. Thus, the deconvolution problem for each bulk tissue sample was represented as a constrained linear least-square problem, which was:

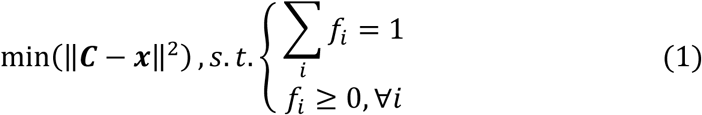

Here, ***f*** is the vector of cell type mixing proportion, and ***C*** is the gene expression matrix of the cell type signature genes in each cell type, while ***x*** is the known expression level of the cell type signature genes. This model has been widely used in the deconvolution problem^23,25^, and can be solved using quadratic programming^62^.

The second method, CIBERSORT^24^, which was developed by Newman et al. in 2015, was also used. Instead of the solving the least square problem, CIBERSORT estimates the relative cell type composition based on nu-support vector regression, minimizing a linear ε-insensitive loss function with L2 regularization.

A third method, namely diffusion ratio (DR) based deconvolution, which is based on the simple assumption that the expression of a cell type signature gene in the bulk tissue can be seen as its expression in the cell type scaled by the cell type’s mixing proportion. It is similar as the recent algorithm MCP-counter ^26^ but was implemented in a simpler way. Under this assumption, the proportion of cell type *i* (represented as *f*_*i*_) was simply estimated as:

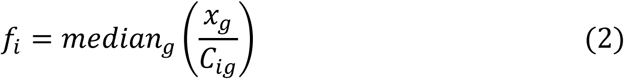

Here, *g* represented each cell type signature gene of cell type *i*. After calculating *f*_*i*_ for all the cell types, ***f*** was normalized so that ∑_*i*_ *f*_*i*_ = 1.

### Composition pattern robustness to confounding factors and cell variability

To investigate the potential influence of confounding factors, including neuronal heterogeneity, neuron ageing, sexes and ethnicity, in transcriptome deconvolution procedure, we acquired genes with differential expression (DE) in different neuron subtypes^57^, neuron ageing^63^ and DE genes between male and female brains^64^ from the previous studies. Genes showing population or sex related different expression level was further obtained using ANCOVA in the age series RNA-seq dataset11 (BH corrected *P*<0.05). The 178 DE genes which were among the cell type signature genes used for DR deconvolution (confounding signature genes) were excluded for the DR transcriptome deconvolution in both healthy and autistic samples. Pearson’s correlation coefficient (PCC) was used to compare the lifespan temporal composition patterns or proportion change patterns in autism before and after exclusion of confounding signature genes.

To investigate the influence of cell variability in the single cell RNA-seq dataset^21^ (Darmanis dataset), we applied 100 bootstrapping of the 420 single cell samples in Darmanis dataset and did DR transcriptome deconvolution based on the cell type gene expressions from bootstrapping. PCC was used to compare bootstrapping-based lifespan temporal composition patterns or proportion change patterns in autism to the patterns based on the full dataset.

### Deconvolution for cell-type expression profile re-estimation

A model similar to the QP-based deconvolution for cell-type composition described above was used for the second deconvolution to re-estimate the cell-type expression profile. Similarly, the deconvolution problem for each gene can be seen as a constrained linear least-square problem, that is:

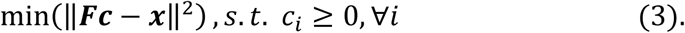

Here, ***c*** was the vector of re-estimated gene expression level in different cell types and ***x*** was the vector of gene expression across the bulk tissue samples. ***F*** was the composition matrix with each row representing the estimated mixing proportion in the corresponding bulk tissue sample. This problem was also solved by using quadratic programming as described above.

### Performance measurement of transcriptome deconvolution

Two different types of deconvolution performance measurements were used to compare different deconvolution methods. The first type of performance measurement, namely similarity measurement, aimed to measure the general similarity between the real bulk tissue gene expression and the predicted gene expression based on the cell type gene expression level and the cell type mixing proportion. It is commonly used to measure deconvolution performance. Here, two measurements were applied: Pearson’s correlation coefficient (PCC), and root-mean-square deviation (RMSD) which was calculated as the root mean squares of log10-transformed fold changes between the observed FPKMs and the predicted FPKMs (products of estimated cell-type proportions and re-estimated cell type expressions) across all genes.

The second type of performance estimation, namely discrimination measurement, was established based on the following assumption: the accurate deconvolution can result in re-estimated cell type expression estimations which are not only similar to the measured cell type expressions, but can also recover the cell type identities discriminating one cell type from the others. Using PCC as the proxy of similarity between two cell type expression profiles, the discriminating recovery score (DRS) of cell type *i* was defined as:

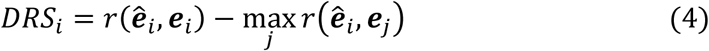

Here, ***e*_*i*_**was the vector of observed log10-transformed FPKM of cell type *i*, and 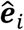 was the vector of re-estimated log10-transformed FPKM of cell type *i*. The mean DRS across different cell types was then used as the proxy of the overall DRS.

### Dissection of composition-dependent and composition-independent expression components

With **X** denoted as the *N × M* gene expression matrix with *N* genes and *M* samples, and the *d × M* estimated composition matrix **F** with *d* as the number of cell types, the re-estimated cell type expression matrix **C** with *N* rows and *d* columns was obtained as described above. The composition-dependent component was then represented as the matrix **X_d_** = **CM**, while the composition-independent component was represented as **X_i_** = **X** - **X_d_**.

### ANCOVA based on natural spline with variable degree of freedom

To identify age-related changes, an ANCOVA employing the F-test was applied to each gene, with its observed expression level, composition-dependent component, or composition-independent component as the response variable. This test, referred as the age test, was used to compare the null model: a linear model only with intercept, to a series of alternative models: the natural spline with degree of freedom from two to eight in response to square root transformed sample ages (sqrt-age). The best alternative model was chosen by applying the adjusted *r*^2^ criterion ^8^. Genes with BH corrected *P*<0.05 were considered as genes with its expression, composition-dependent component or composition-independent component changed in the age-related manner. For calculating the proportion of variance explained by age, the natural spline model with fixed degree of freedom (*df*=8) in response to sqrt-age was used.

The similar ANCOVA framework was also applied to identify changes between autistic samples and healthy samples, by considering the sample ages. The null model was defined as the unified natural spline model with no discrimination of autistic and healthy samples, while the alternative model with varied coefficient in the two groups was compared by employing the F test. Genes with BH corrected *P*<0.2 were considered as genes with composition-dependent or composition-independent component changed between the two groups of samples. Similarly, sqrt-age was used as the independent variable.

### Cell type expression enrichment analysis

The cell type enrichment analysis was performed based on the average RPKM of each gene in the eight major cell types in human brains including replicating neurons, immature neurons, mature neurons, astrocytes, oligodendrocytes, endothelial cells, microglia and oligodendrocyte precursor cells (OPCs), based on the human brain single cell RNA-seq data^21^. The expression specificity of one gene in one cell type was represented by the log10-transformed fold change between its expression level in the cell type and its average expression level in the other cell types. For each cell type, a one-sided Wilcoxon’s rank test was applied to compare the expression specificity of genes in the gene list, to the other detected genes.

## Acknowledgement

We thank Prof. Dr. P. Khaitovich, Dr. Y. Wei, Dr. S. Feng, Dr. H. Hu, Ms. H. Bammann, Ms. Q. Li, and Mr. C. Xu for the valuable discussions. We thank Dr. G. L. Banes for his helpful comments on the manuscript.

## Author Contributions Statement

Q.Y. and Z.H. conceived and designed the study, analyzed the data, and wrote the manuscript. Both the authors have read and approved the final manuscript.

## Data availability

The fetal and adult human brain single cell RNA-seq for cell type gene expression reference is available in GEO (accession: GSE67835). The postnatal human brain age series RNA-seq data is available in GEO (accession: GSE51264). The autistic human brain RNA-seq data is available in GEO (accession: GSE59288).

